# Overfeeding does not increase energy expenditure or energy excretion in mice

**DOI:** 10.1101/2024.04.06.588219

**Authors:** Pablo Ranea-Robles, Camilla Lund, Jens Lund, Maximilian Kleinert, Christoffer Clemmensen

## Abstract

To curb the obesity epidemic, it is imperative that we improve our understanding of the mechanisms controlling fat mass and body weight regulation. While great progress has been made in mapping the biological feedback forces opposing weight loss, the mechanisms countering weight gain remain less well defined. Here, we integrate a mouse model of intragastric overfeeding with a comprehensive evaluation of the regulatory aspects of energy balance, encompassing food intake, energy expenditure, and fecal energy excretion. To evaluate the role of adipose tissue thermogenesis in the homeostatic protection against overfeeding-induced weight gain, we exposed uncoupling protein 1 (UCP1) knockout (KO) mice to overfeeding. Our results confirm that 7 days of 150% overfeeding induces ∼11% weight gain and triggers a potent and prolonged reduction in voluntary food intake that drives body weight back to baseline following overfeeding. Overfeeding has no effects on energy expenditure, consistent with the observation that mice lacking UCP1 are not compromised in their ability to defend against overfeeding-induced weight gain. These data emphasize that whole-body energy expenditure and adipose thermogenesis are not key contributors to protection against overfeeding in mice. Lastly, we show that fecal energy excretion decreases in response to overfeeding, primarily driven by a reduction in fecal output rather than in fecal caloric content. In conclusion, these results challenge the prevailing notion that adaptive thermogenesis contributes to the defense against weight gain induced by overfeeding. Instead, the protection against enforced weight gain in mice is primarily linked to a profound reduction in food intake.

## Introduction

Obesity is a disease with multifactorial etiology that poses a significant risk for a series of severe comorbidities including type 2 diabetes, cardiovascular diseases, and cancers (1). Despite considerable progress in developing effective weight loss therapies, long-term weight management strategies remain elusive, and the underlying causes of obesity are a subject of intense debate (2). Anti-obesity interventions typically result in rapid weight loss followed by a weight plateau and progressive regain. This weight regain is thought to be triggered by the body perceiving weight loss as a threat to homeostasis, prompting a response characterized by hypoleptinemia, increased appetite and decreased energy expenditure (3). Mirroring this defensive response, the organism similarly perceives deliberate attempts to gain weight as a homeostatic insult and engages compensatory responses to restore energy balance (2,4). However, the mechanisms that counteract positive energy balance and weight gain remain less understood (2,4–7).

For decades, it has been debated whether overfeeding-induced weight gain is countered by so-called *luxuskonsumption*, an adaptive increase in energy expenditure beyond what can be attributed to an increased body mass (3,4,8–10). Some rodent studies using intragastric overfeeding and indirect metabolic measures have observed subtle (11) or transient (6) increases in whole-body energy expenditure, linked to thyroid hormones (12) and norepinephrine (11,13). However, other studies have not detected changes in energy expenditure during (14–16) or after (17) overfeeding (for review, see 8). This inconsistency highlights the need for clearer evidence on whether adaptive thermogenesis consistently helps counteract weight gain after overfeeding. Similarly, the role of fecal energy excretion in body weight regulation remains insufficiently characterized (18). In healthy humans, between 1 and 11% of ingested energy appears to be lost through stool (19) and large interindividual variability in fecal calorie loss has been observed in the context of overfeeding (ranging from 80 to 500 kcal/day) (19). This highlights the potential impact of fecal energy excretion on body weight, as losing such a substantial amount of energy will reduce the amount of metabolizable energy.

In this study, we employed a mouse model of intragastric overfeeding (20) combined with highresolution indirect calorimetry in metabolic cages and bomb calorimetry of fecal outputs to comprehensively assess temporal changes in energy intake, whole-body expenditure, and fecal energy excretion in response to overfeeding. Additionally, to further explore the role of adipose thermogenesis in the homeostatic response to overfeeding, we subjected uncoupling protein 1 (UCP1) knockout (KO) mice to intragastric overfeeding.

## Results

### Overfeeding-induced changes in body weight and energy intake

We measured changes in body weight and voluntary food intake in mice subjected to 7 days of intragastric overfeeding (OF) (150%, i.e. 50% excess calories) followed by 4 days of recovery (Rec) (Figure 1A). Overfed mice gained 10.7% weight on average (Figure 1B), corresponding to 3.4 g (Figure 1C), whereas control mice remained weight stable (Figure 1B,C). After overfeeding, mice returned to their original body weight within 4 days (Figure 1B,C). Despite similar body weights in control and overfed mice after 4 days of recovery, overfed mice exhibited a slightly higher percentage of fat mass and a slightly lower percentage of lean mass compared to controls (Figure 1D). Consistent with our previous results as well as other reports (5,6,11,12,14,17,20,21), weight gain during overfeeding occurred together with a marked reduction in voluntary food intake which gradually returned to baseline levels after calorie infusion stopped (Figure 1E). Overfeeding was also associated with a pronounced reduction in water intake that persisted throughout the recovery period (Figure 1F).

**Figure 1.**
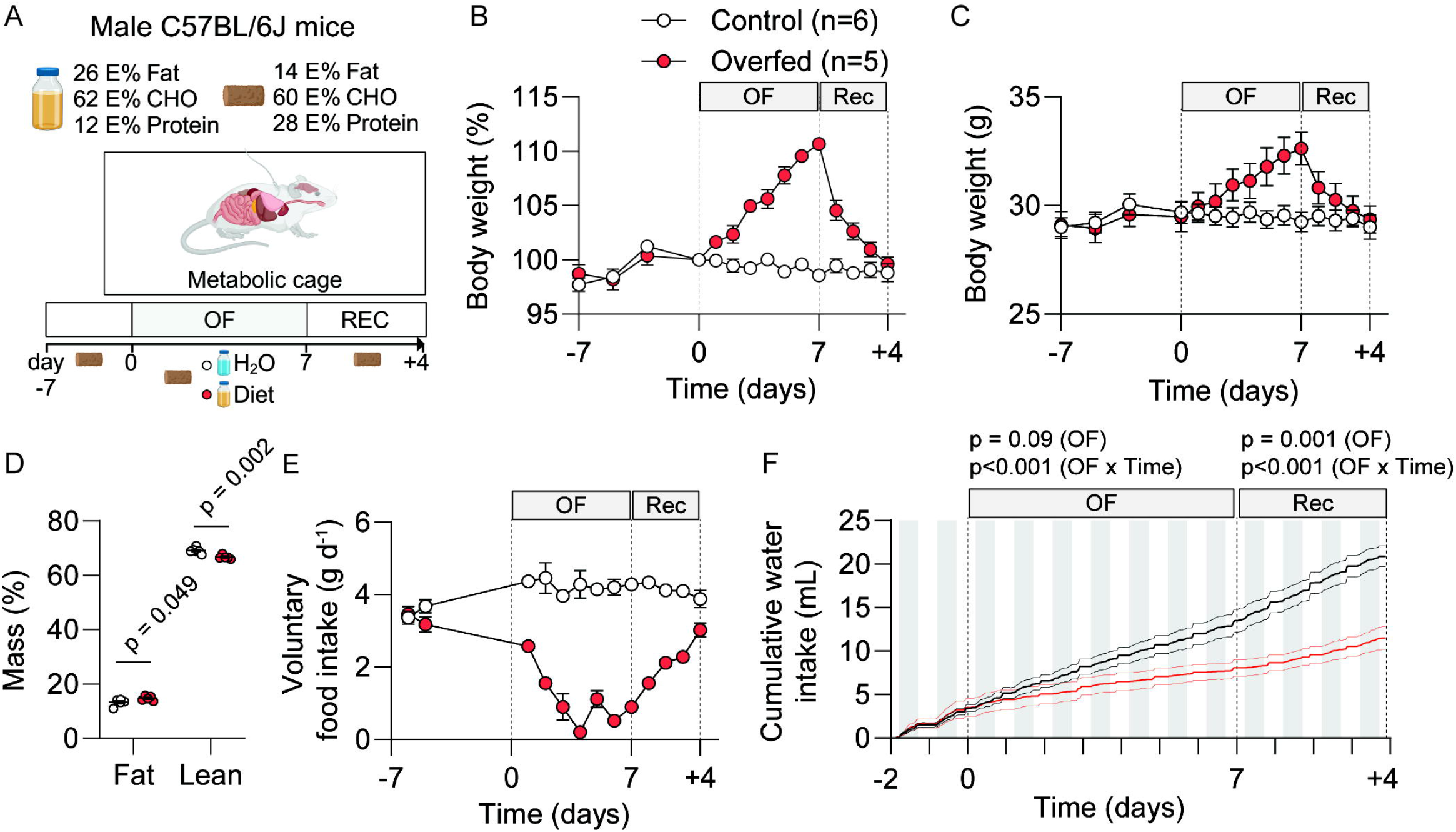
Overfeeding-induced changes in body weight and energy intake. **A**) Schematic overview of the experimental overfeeding setup in indirect calorimetry cages. **B**) Body weight changes (%,100% at day 0) in control (n=6) and overfed (n=5) mice. **C**) Absolute body weight changes (in grams). **D**) Body composition (percentage of fat and lean mass) at the end of the recovery period (d+4, postmortem). **E**) Voluntary food intake (in grams). **F**) Cumulative water intake (in mL). Data shown as mean ± SEM (B-F) with individual values plotted in D. All data correspond to the same set of control (n=6) and overfed (n=5) mice. Overfeeding (OF) and recovery (Rec) periods are indicated by vertical dashed lines. p values were calculated using multiple unpaired t tests with Welch correction after adjusting for multiple comparisons with a False Discovery Rate set at 1% (D), or 2way ANOVA (F) using overfeeding and time as factors.

### Energy expenditure and UCP1-mediated adipose thermogenesis in response to overfeeding

We found no difference in average daily energy expenditure between the groups, both during overfeeding and recovery periods (Figure 2A,B). Although mice displayed the expected circadian variations with a higher metabolic rate during their active phase, overfeeding itself did not induce an overall increase in energy expenditure (Figure 2A). This finding was confirmed by regression-based ANCOVA controlling for body weight (Figure 2B). These results demonstrate that 7 days of overfeeding do not cause an adaptive increase in energy expenditure. In contrast, overfeeding triggered significant shifts in metabolic fuel utilization, revealed by changes in the respiratory exchange ratio (RER). During overfeeding, mice relied more on carbohydrates, indicated by an elevated RER (Figure 2C,D). This shifted rapidly after infusion stopped, with RER dropping sharply in overfed mice (Figure 2C,D) and gradually returning to control levels by day 4 of recovery (Figure 2C), reflecting a swift change to fat oxidation after overfeeding. While previous studies have linked increased non-exercise activity thermogenesis (NEAT) to weight gain resistance in humans (22), we did not observe any significant differences in locomotor activity during overfeeding or recovery periods (Figure 2E,F). However, a trend towards decreased locomotor activity was observed in the light phase during the overfeeding period (Figure 2F).

**Figure 2.**
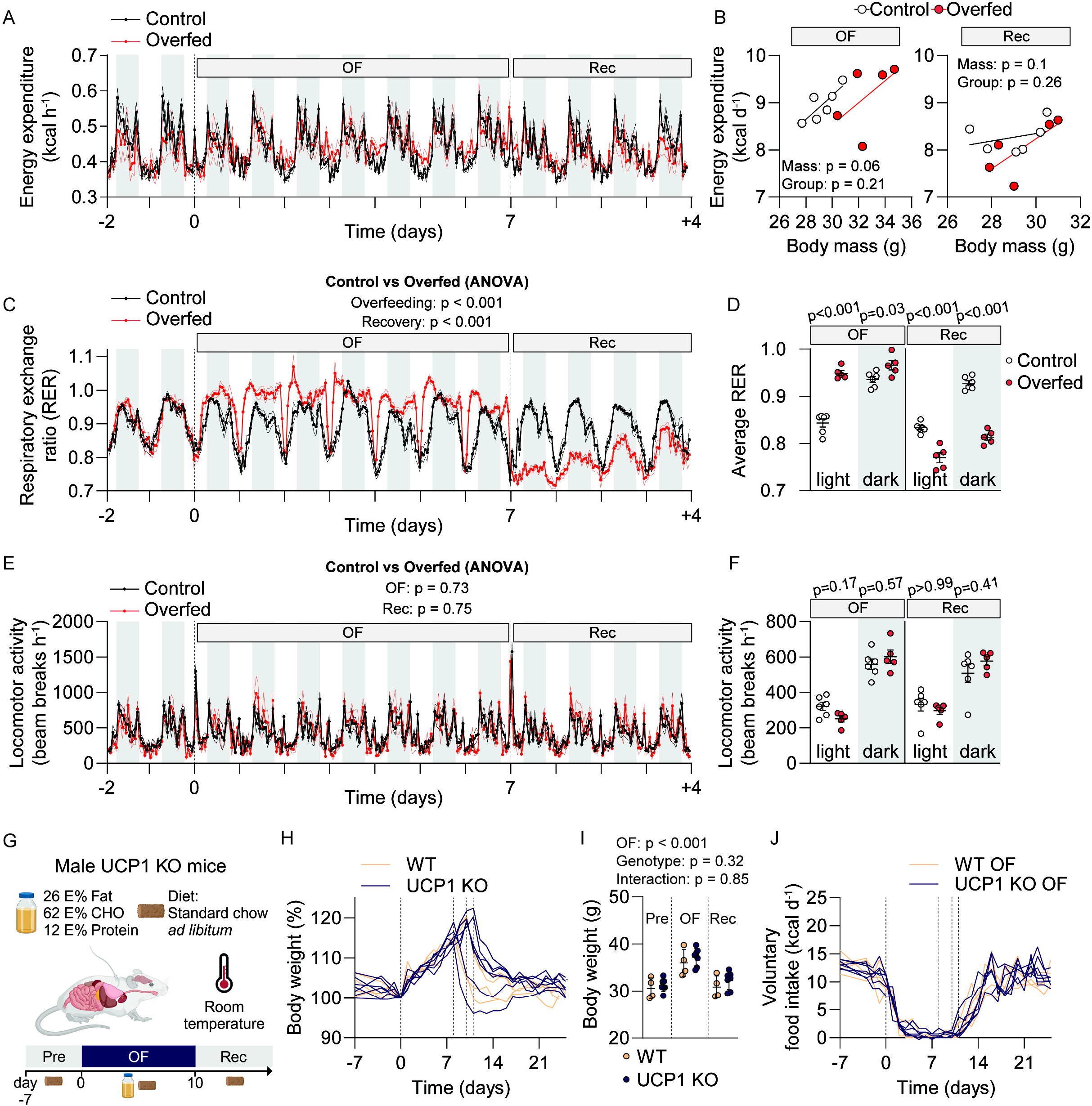
Energy expenditure and UCP1-mediated adipose thermogenesis in the response to overfeeding. **A**) Whole-body energy expenditure (kcal per hour) in control (n=6) and overfed mice (n=5) from Figure 1. **B**) Regression-based ANCOVA using body mass as a covariate during the overfeeding period (OF) and recovery (Rec) periods. **C**) Respiratory exchange ratio (RER) during the overfeeding period (OF) and recovery (Rec) periods. **D**) Average RER during light and dark phases in the overfeeding and recovery periods. **E**) Locomotor activity (XYZ beam breaks per hour). The transient surge in locomotor activity during day 0 and 7 reflect cage changes with provision of clean bedding. **F**) Average locomotor activity (beam breaks per hour) during light and dark phases in the overfeeding and recovery periods. **G**) Schematic overview of the experimental overfeeding setup in germline UCP1 KO mice. **H**) Body weight changes (%,100% at day 0) in WT (n=4) and UCP1 KO (n=7) overfed mice. Individual trajectories are shown (mice finished OF on d8, d10 or d11, indicated by vertical dashed lines). **I**) Body weight (in grams) of the same mice as shown in H. **J**) Voluntary food intake (kcal per day) of same mice shown in H-I. Data shown as mean ± SEM (A,C-F,I). Individual data points (B, D, F, I) or individual trajectories (H,J) are shown. Night phase shown as light grey shades (A, C-F). p values were calculated using ANCOVA with body mass as a covariate (B), ANOVA from CalR (C, E), and 2-way ANOVA (D,F,I). p values are indicated. p values in D and F represent post-hoc comparisons after two-way ANOVA in each part of the day (light and dark phases) using overfeeding (control and overfed) and period (overfeeding and recovery) as factors. OF: Overfeeding. Rec: Recovery.

We also investigated the potential role of adipose thermogenesis in the response to overfeeding, which might not be detectable by our energy expenditure measurements. UCP1, a protein found in the inner mitochondrial membrane, uncouples substrate metabolism from ATP synthesis (23), thus generating heat and potentially counteracting overfeeding-induced weight gain. We subjected wild-type (WT) and germline UCP1 KO mice to overfeeding (Figure 2G) and found similar relative (+17.7% in WT vs. +19% in UCP1 KO mice) and absolute weight changes (+5.4 g in WT vs. +6.0 g in UCP1 KO mice) (Figure 2H,I). Weight loss after overfeeding was rapid and genotype-independent (Figure 2H,I). Additionally, WT and UCP1 KO mice exhibited similar suppression in voluntary food intake during overfeeding, gradually returning to baseline levels after overfeeding (Figure 2J). These findings suggest that UCP1-mediated adipose thermogenesis is not essential for the protective response to experimental overfeeding in mice housed at standard room temperature (22°C).

### Changes in fecal energy excretion and energy balance in response to overfeeding

To measure fecal energy excretion, we measured fecal energy content using bomb calorimetry. While fecal energy density remained unchanged by overfeeding (Figure 3A), overfed mice excreted around 3 times less feces during both overfeeding and recovery periods (Figure 3B), leading to lower fecal energy excretion (Figure 3C). This reduction possibly reflects increased absorption of the liquid overfeeding diet compared to solid fiber-rich chow, as suggested by the increased digestive efficiency observed in overfed mice (95%) compared with control mice (81%) (Figure 3D). Energy balance calculations revealed a positive energy balance in overfed mice during overfeeding followed by a negative energy balance during recovery, as expected (Figure 3E), whereas control mice displayed a slightly positive energy balance in both phases (Figure 3E). Overfed mice exhibited a higher positive energy balance across the entire study compared to controls, despite similar body weights (Figure 3E). These findings suggest that while fecal energy excretion plays a minor role in the homeostatic response to overfeeding, the higher digestive efficiency in the recovery phase might contribute to higher positive energy balance after four days of weight recovery.

**Figure 3.**
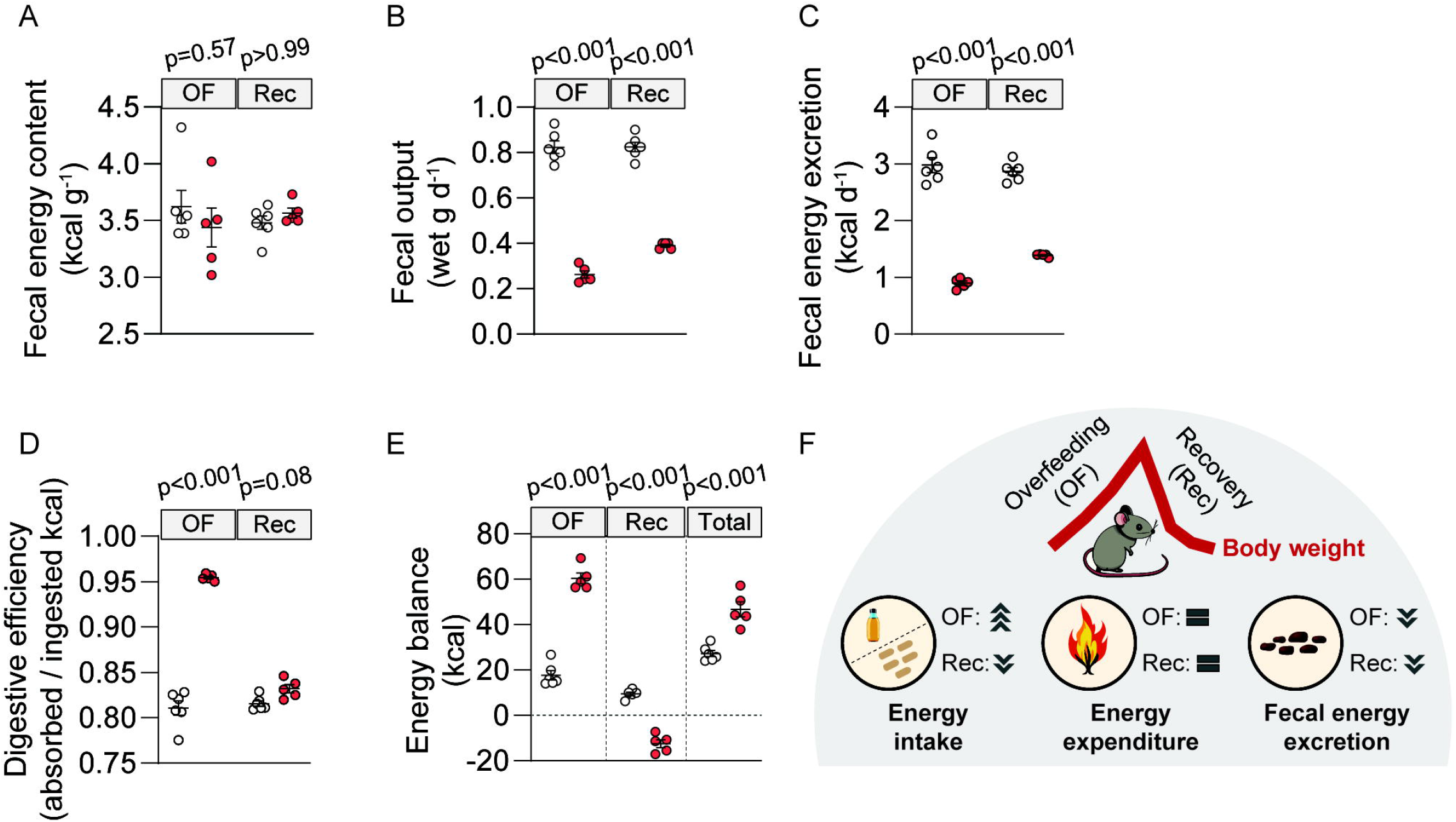
Changes in fecal energy excretion and energy balance in response to overfeeding. **A**)Fecal energy content (kcal per gram) measured by bomb calorimetry at the end of the OF and Rec periods in control (n=6) and overfed (n=5) mice from Figure 1. **B**) Fecal output (wet grams per day) during OF and Rec periods. **C**) Daily fecal energy excretion (kcal per day) during OF and Rec periods. **D**) Digestive efficiency (energy absorbed / energy ingested) at the end of the OF and Rec periods. **E**) Estimated energy balance at the end of the OF, Rec, and combined (total) periods. **F**) Schematic figure showing the changes in body weight and energy balance components (energy intake, energy expenditure and fecal energy excretion) during overfeeding and recovery. Data shown as means ± SEM with individual values plotted. p values were calculated using 2-way ANOVA and Bonferroni’s post-hoc analysis using overfeeding and period as factors. p values are indicated. OF: Overfeeding. Rec: Recovery.

## Discussion

The mechanisms by which energy balance and body weight are regulated in response to overfeeding remain unclear. Here, we demonstrate that energy expenditure does not increase with 7 days of intragastric overfeeding in mice. Furthermore, we observe a decline in fecal energy excretion following overfeeding, predominantly attributed to decreased fecal output rather than a reduction in fecal caloric content. Our findings suggest that the rapid return to normal body weight upon cessation of overfeeding is predominantly due to decreased food intake rather than increased adaptive thermogenesis or fecal energy loss.

Rodent studies investigating changes in energy expenditure during overfeeding present contrasting findings. Two rat studies observed a transient increase in energy expenditure at the end of overfeeding periods, with the effects disappearing by either day 1 (6) or day 3 (11) into the recovery phase. Conversely, other studies in rats (14–16) found no changes in energy expenditure beyond those attributable to increased body size during overfeeding, even at high calorie surpluses (200%). Similarly, the only mouse study (to our knowledge) measuring energy expenditure in response to overfeeding showed no increase after two weeks of intragastric overfeeding (17). While Ravussin et al. have suggested that a decrease in energy efficiency (weight gain per calorie ingested) following overfeeding in mice could indicate increased energy expenditure or excretion (17), our current findings, based on the integration of indirect calorimetry and intragastric overfeeding, suggest otherwise. It remains to be determined if prolonged intragastric overfeeding increases energy expenditure and if other variables, such as varying levels of caloric excess and differences in animal species and strains, affect energy expenditure.

Assessment of energy expenditure in human overfeeding trials is challenging and has yielded highly variable and inconsistent results (4,8). Our findings in mice, showing no significant increase in energy expenditure alongside similar weight gain and recovery patterns in UCP1 KO mice compared to wildtype littermates in response to overfeeding, align with some previous human studies suggesting minimal brown adipose tissue activation and limited changes in energy expenditure due to overfeeding (8–10,24). This supports the notion that luxuskonsumption might not be a major contributor to the defense against weight gain in humans. However, some human studies have found an increase in total energy expenditure in response to overfeeding (3,25). These contrasting findings and methodological differences between human studies and rodent intragastric overfeeding limit the direct translation of our mouse data to humans. While our study underscores the limited impact of energy expenditure and UCP1-mediated thermogenesis in mitigating overfeeding-induced weight gain in mice, this does not preclude the possibility of adaptive increases in energy expenditure and/or energy excretion offering protection against weight gain in other scenarios. For instance, some mouse strains and humans with higher resistance to obesity may exhibit such adaptive responses (4,18). In addition to factors such as overfeeding duration, calorie excess, and interindividual variability, the potential influence of diet composition on the response to overfeeding should also be considered. Unlike our study, which utilized a liquid diet lacking fiber, human overfeeding studies typically involve solid foods. Fiber is known to influence fecal energy excretion and energy balance (26). Human studies have reported an increase in total fecal energy output in response to overfeeding (3,19), potentially explained by the presence of fiber in the diet, which promotes fecal bulking and potentially higher energy excretion compared to a fiber-deficient liquid diet.

In this study, we integrate intragastric overfeeding in rodents with high-resolution assessments of whole-body energy expenditure, metabolic fuel utilization, and locomotor activity across both overfeeding and recovery phases. This comprehensive approach, complemented by measurements of fecal energy excretion, provides a detailed picture of the metabolic and energy balance changes associated with short-term overfeeding in mice. One of our primary findings - that adjustments in food intake serve as a primary adaptive response to intragastric overfeeding - aligns with prior research indicating significant appetite suppression in animals subjected to overfeeding (5,20,21,27). While we found no evidence of overfeeding-induced thermogenesis, the immediate shift towards fat oxidation post-overfeeding might constitute an important adaptive response contributing to resistance against excessive weight gain (28). Hence, while rapid changes in food intake seem to predominantly drive alterations in body weight during overfeeding, early shifts in metabolic fuel oxidation may causally contribute to the hypophagic response (29). Further enhancing temporal data resolution could be valuable in elucidating the precise sequence of physiological changes underlying the homeostatic response to overfeeding. Additionally, longer overfeeding interventions, employing larger animal models, exploring diverse diet compositions, and evaluating alternative routes of energy loss such as urine and skin (18,30), would offer a more comprehensive understanding of the physiological response to overfeeding.

## Methods

### Animal husbandry

All mouse studies were conducted at the University of Copenhagen, Denmark, and carried out in accordance with regulations regarding the care and use of experimental animals that were approved by the Danish Animal Experimentation Inspectorate (2018-15-0201-01457; 2023-15-0201-01442). Wild type (WT) male C57BL/6J mice (Janvier, FR), and germline UCP1 whole-body KO mice (31) (*Ucp1*^*tm1Kz*^, The Jackson Laboratory, stock #003124) were used. All experiments were done at 22□C with a 12:12 h light-dark cycle (6am-6pm). Mice had ad libitum access to water and chow diet (SAFE D30, Safe Diets, France). All mice were single housed after surgery and during experimental overfeeding and recovery.

### Experimental overfeeding

The surgical procedure to insert the gastric catheter and the material used for automated overfeeding were the same as previously described (20). The methodological aspects of the studies conducted here are described in detail in the Supplementary Methods.

### Fecal energy excretion

For assessment of excreted energy via feces (32), fecal pellets were collected after the overfeeding (7 days) and recovery (4 days) periods, respectively. The bedding and nesting content of every cage was dried at room temperature for 1 week. Then, the cage bedding was separated by size using kitchen sieves with varying mesh sizes (Veras Verden, Denmark). The flow through was collected in a tray and manually examined for food remnants and fecal pellets. Food remnants were weighed to determine total food spillage and this was adjusted for in the calculation of food intake for energy balance in Figure 3. Feces were carefully collected using tweezers, weighed, and desiccated in a drying oven (50□°C) before bomb calorimetric combustion (IKA C5003, IKA Werke, Germany). The absorbed energy was determined by combusting a representative sample of chow diet calorimeter. The digestive efficiency was calculated by dividing the absorbed energy (energy intake – energy lost with feces) by the ingested energy.

### Data analysis

Data were analysed and plots were generated using GraphPad Prism software version 10.1. All data are presented as mean□±□SEM, unless indicated otherwise. Findings with p values□≤□0.05 were considered statistically significant. All statistical analyses are indicated in figures and figure legends. Indirect calorimetry data were exported with Macro Interpreter, macro 13 (Sable Systems) prior to analysis in CalR version 1.3 (33). Regression-based ANCOVA analysis of energy expenditure using body weight as a covariate were performed using absolute body weight at d7 and d7+4 for the overfeeding and recovery period, respectively.

## Acknowledgments

We would like to thank Morten Dall and members of the Rodent Metabolic Phenotyping Platform at the Novo Nordisk Foundation Center for Basic Metabolic Research for their help with intragastric surgeries, data export and setup of the Sable System. We also thank members of the Clemmensen laboratory for stimulating discussions and Cláudia Gil, Charlotte Svendsen, Vaida Juozaityté for helping with collection of fecal pellets. We thank Johanna Bruder (Martin Klingenspor group, Technical University of Munich) for kindly providing advice on fecal pellet collection. Tao Ma and Astrid Linde Basse (Gerhart-Hines group at the CBMR) kindly provided founders to establish a UCP1 mouse colony for overfeeding experiments. JL is supported by the BRIDGE – Translational Excellence Programme (www.bridge.ku.dk) at the Faculty of Health and Medical Sciences, University of Copenhagen, funded by the Novo Nordisk Foundation (NNF20SA0064340). Financial support to CC was provided through Lundbeck Foundation (Fellowship R238-2016-2859) and the Novo Nordisk Foundation (Grant number NNF17OC0026114). The Novo Nordisk Foundation Center for Basic Metabolic Research is an independent Research Center, based at the University of Copenhagen, Denmark, and partially funded by an unconditional donation from the Novo Nordisk Foundation (www.cbmr.ku.dk) (Grant number NNF18CC0034900). M.K. was supported by the Deutsche Forschungsgemeinschaft (DFG; KL 3285/5-1), the German Center for Diabetes Research (DZD; 82DZD03D03 and 82DZD03D1Y), and the Novo Nordisk Foundation (NNF; NNF19OC0055192).

## Author contributions

PRR, CL, and CC conceived the study. PRR & CL performed animal surgeries and executed the mouse *in vivo* studies. MK analyzed the energy content of food and feces. PRR analyzed the data and wrote the draft of the manuscript. CL & JL reviewed and edited the draft of the manuscript. CC supervised the study, reviewed, and edited the manuscript. All co-authors contributed to data interpretation and provided input to the manuscript.

## Competing interests

CC is a co-founder of Ousia Pharma ApS, a biotech company developing therapeutics for treatment of metabolic disease. The remaining authors declare no competing interests.

