## Supplementary Methods for "Overfeeding does not increase energy expenditure or energy excretion in mice"

**Experimental overfeeding**

*Study 1: Experimental overfeeding in metabolic cages*

Following a modified surgical protocol (1,2) that we have previously described (3), 18-week-old male C57BL/6J (n=16) were anesthetized with isoflurane and subjected to a midline abdominal incision to expose the gastric ventricle. A purse-string suture was placed above the gastric fundus, and a 22-gauge catheter (C30PU-MGA2209, Instech Laboratories, USA) was inserted through the center of the suture into the lumen of the stomach. A subcutaneous vascular access button (VABM1B/22, Instech) was secured to the skin at the back of the mouse neck, and the catheter was tunneled under the skin to connect the button. After surgery, mice were single housed and allowed to recover for 5 weeks before proceeding with overfeeding interventions. After recovery from surgery, individual body weight and food intake were tracked for 3 days before placing mice in metabolic cages. Mice were then divided into two groups, control (n=6) and overfed (n=10), matched by absolute body weight at the day before study start. Prior to their placement into an indirect calorimetry system, mice were connected to the automated infusion system. This system consists of infusion pumps (704500, 704501, 703005, 703024, Harvard apparatus, USA) and syringes (10 mL syringes with Luer Lock tip, 302995, Becton and Dickinson, USA) connected to the vascular access buttons using 22ga tethers with springs (VABM1T/22, Instech Laboratories, USA), polyethylene tubing (BTPE-50, Instech Laboratories, USA), multi-axis lever arms (SMCLA, Instech Laboratories, USA), and 22ga swivels (375/22PS, Instech Laboratories, USA). Mice always had *ad libitum* access to chow and tap water during the entire study. Mice movement was not limited due to being connected to the infusion system.

The study was initiated after 5 days of acclimatization in metabolic cages (Promethion Core Metabolic System, Sable Systems International). All mice were continuously infused with sterile water (100 µL/hr) during the last 3 days of this acclimatization period. Then, on day 0, the overfeeding group received 1 day of eucaloric liquid diet (100%) infusion and 6 days of 50% caloric surplus over baseline requirements (150%), whereas control mice were infused with the same volume of sterile water. Every day at 10 am during the overfeeding period, the infusion system was stopped for 30 minutes, enabling replacement of the syringe with fresh liquid diet and flushing of the outer tubing with sterile water. After syringe replacement, mice received an injection of 0.5 mL sterile water through the vascular access button and were infused with 1000 µL/h sterile water for 30 minutes (another 0.5 mL). Therefore, the amount of diet needed to achieve 150% of overfeeding was calculated for 23 hours of infusion per day. After 7 days of overfeeding, mice remained connected to the infusion system and were allowed to recover for 4 days while receiving 100 µL/h of water infusion. At the end of the experiment, mice were disconnected from the automated overfeeding system and body composition was measured by magnetic resonance imaging (Bruker LF90II). The overfeeding group was infused with a commercial liquid diet (584421, Nutridrink Vanilla, Nutricia, Netherlands) that was supplemented with 12.5% (w/v) sucrose (S0389, Sigma-Aldrich, USA). The addition of sucrose increased the caloric content of the liquid diet from 1.5 kcal/mL to 2 kcal/mL (fats 26 E%, Carbohydrates 62 E%, protein 12E%). To calculate the flow rate of diet infusion, the daily food intake of the chow (estimated caloric density = 3.389 kcal/g) during the baseline period was averaged for all the mice and divided by the caloric density of the diet (2 kcal/mL). Specific flow rates, daily volume infused, and daily caloric infusion are indicated in Supplementary Table 1. 5 mice in the overfeeding group were excluded from the study due to tube clogging during the overfeeding period that impeded the completion of the diet infusions. Due to imprecisions in the body weight and food intake measurement with the automated system incorporated in the metabolic cages, we measured them manually daily, when syringes were replaced with fresh liquid diet. We also corrected for food spillage at the end of the overfeeding and recovery period, respectively, as indicated in the fecal energy excretion subsection in Methods. Metabolic and behavioural parameters were measured every 15 min, and data was binned and averaged per hour before plotting them into graphs (Supplementary Table 2).

*Study 2: Experimental overfeeding of WT and UCP1 KO mice*

Chow-fed male WT (n=4) and UCP1 KO (n=7) mice on a C57BL/6J background and at 22 to 30 weeks of age were overfed for 10 days as described above with the minor modification that the infusion flow rate was increased gradually until reaching 150% of energy infusion on day 9 (Supplementary Table 1). Mice were matched for baseline body weight (WT = 30.6 ± 2.2 g, UCP1 KO = 31.3 ± 1.4 g) and daily food intake (WT = 13.1 ± 1.8 kcal, UCP1 KO = 13.2 ± 0.7 kcal). Mice were observed after overfeeding to evaluate the changes in body weight and food intake until they stabilized.
